# Heritability estimation of cognitive phenotypes in the ABCD Study^®^ using mixed models

**DOI:** 10.1101/2022.10.28.512918

**Authors:** Diana M. Smith, Robert Loughnan, Naomi P. Friedman, Pravesh Parekh, Oleksandr Frei, Wesley K. Thompson, Ole A. Andreassen, Michael Neale, Terry L. Jernigan, Anders M. Dale

## Abstract

Twin and family studies have historically aimed to partition phenotypic variance into components corresponding to additive genetic effects (*A*), common environment (*C*), and unique environment (*E*). Here we present the ACE Model and several extensions in the Adolescent Brain Cognitive Development Study (ABCD Study^®^), employed using the new Fast Efficient Mixed Effects Analysis (FEMA) package. In the twin sub-sample (*n* = 924; 462 twin pairs), heritability estimates were similar to those reported by prior studies for height (twin heritability = 0.86) and cognition (twin heritability between 0.00 and 0.61), respectively. Incorporating SNP-derived genetic relatedness and using the full ABCD Study^®^ sample (*n* = 9,742) led to narrower confidence intervals for all parameter estimates. By leveraging the sparse clustering method used by FEMA to handle genetic relatedness only for participants within families, we were able to take advantage of the diverse distribution of genetic relatedness within the ABCD Study^®^ sample.

## Introduction

For over a century, researchers have relied on variance partitioning as a statistical method for estimating heritability (Carey 2003). Historically, twin studies provided an avenue by which researchers could model the variance of a given phenotype as comprised of distinct components: e.g., additive genetic effects (*A*), common environmental effects (*C*), and unique environmental effects (*E*, also including error or unmodeled unexplained variance; Martin and Eaves 1977; Neale and Maes 2004). Specifically, in a linear mixed-effect (LME) regression model,

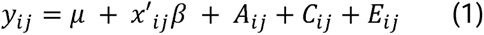

where *y_ij_* is the trait value of individual *j* in family *i*; μ is the overall mean; *x_ij_* denotes a vector of covariates; and *A_ij_*, *C_ij_*, *E_ij_* represent latent additive genetic, common environmental and unique environmental random effects (ACE model), respectively. For longitudinal datasets in which participants are followed over time, the LME is also the recommended analysis that takes into account the random effect of subject (Pinheiro 2014), e.g.:

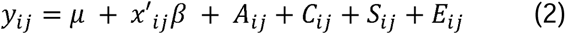

where *S_ij_* is the random effect of subject (e.g., subject ID), and the vector of covariates *x_ij_* typically includes a fixed effect of time (e.g., age). In this way, the longitudinal LME assumes a linear combination of fixed and random effects to model the trajectory of the phenotype of interest over time.

Although the ACE model has often been implemented using structural equation model (SEM) software such as *OpenMx* (Neale et al. 2016), the SEM representation is mathematically equivalent to the LME regression model shown in Equation 1 (Neale and Maes 2004; Visscher et al. 2004; McArdle and Prescott 2005). Indeed, prior applications of the ACE framework have been implemented using LMEs from R and Stata packages (Rabe-Hesketh et al. 2008) as well as SAS (Wang et al. 2011). For studies that incorporate extended family designs with several random effects, Visscher and colleagues (2004) recommended implementation using a LME approach, though SEM methods also exist to model complex family structure (Truett et al. 1994; Keller et al. 2009).

With recent advances in genomic sequencing, there has been an influx of methods that use measured genetic data rather than inferred genetic similarity from twin status. For example, genome-wide complex trait analysis (GCTA; Yang et al. 2011) was developed to incorporate a pairwise genetic relatedness matrix (GRM) between individuals using information from single nucleotide polymorphisms (SNPs). Twin studies have subsequently been adapted to incorporate empirical measures of genetic relatedness (Kirkpatrick et al. 2021). However, incorporating a matrix of pairwise relatedness values for each set of participants leads to an increase in the computational time when estimating these model parameters. Various subsequent adaptations have been developed to increase the processing speed of GCTA software (Ge et al. 2015) and to incorporate effects of maternal and/or paternal genotype on the traits within GCTA (Eaves et al. 2014; Qiao et al. 2020; Eilertsen et al. 2021).

Comparison of heritability estimates derived from non-twin versus twin analyses have found that non-twin studies consistently yield lower heritability estimates, an example of the so-called “missing heritability” in genetics research (Kim et al. 2015). Some researchers have suggested that this phenomenon may be due, in part, to inflated twin heritability estimates; for example, due to dominant genetic variation which might be masked by shared environment in twin and family studies (Chen et al. 2015). Indeed, twin and family studies have developed several ways of parsing “common environment”, including using geospatial location information (Heckerman et al. 2016; Fan et al. 2018) and adding a random effect of twin status (T) when including twins and full siblings in the same study (Zyphur et al. 2013). It should also be noted that “SNP heritability” is a “narrow heritability” that only takes into account the additive genetic effects, and can be an underestimation of true heritability depending on SNP coverage. For example, estimates of SNP heritability tend to increase as SNP coverage increases from 300,000 SNPs to whole genome sequencing, and as rarer genetic variants are included (Wainschtein et al. 2022).

The Adolescent Brain Cognitive Development Study (ABCD Study^®^) provides a particularly appealing dataset for the estimation of heritability, not only due to its population sampling framework, large sample size, and longitudinal design, but also because it contains an embedded sub-sample of 840 pairs of same-sex twins recruited through birth registries at four sites (Iacono et al. 2018). The overall sample is thus enriched for genetic relatedness, with families that include siblings, half siblings, dizygotic (DZ) twins, and monozygotic (MZ) twins. The ABCD Study^®^ data therefore requires the application of modeling approaches that take repeated measures, family structure, and relatedness into account.

In this study we implemented modeling strategies that account for family structure and pairwise genetic relatedness using the recently developed Fast Efficient Mixed Effects Analysis (FEMA; Fan et al. 2021). We used FEMA to model participants nested within families, where random effects such as genetic relatedness were taken into account for each pair of subjects within a family, and set to zero for individuals who are not in the same family (Fan et al. 2021). For longitudinal data FEMA also allows inclusion of the random effect of subjects, which is a common practice for repeated-measures designs (Pinheiro 2014). FEMA provides a flexible platform for users to specify a wide array of fixed and random effects, which makes it a useful tool for modeling variance components in the ABCD Study^®^. The ABCD Study^®^ Data Analysis, Informatics & Resource Center (DAIRC) intends to incorporate FEMA into the Data Exploration and Analysis Portal (DEAP) so that investigators can easily specify and run LMEs through this online platform. This paper is therefore intended to serve as a reference point for users who are examining random effects estimates on behavioral phenotypes using FEMA.

We first compared the basic ACE model implemented in FEMA versus *OpenMx* (Neale et al. 2016). Next, we tested the effect of including SNP-derived genetic relatedness (using genotype array data) compared to estimating relatedness based on kinship (i.e., 1.0 for MZ twins, 0.5 for DZ twins and full siblings). We then progressively expanded our sample size, first going from “twins only” to the full ABCD Study^®^ baseline sample (including non-twin siblings and singletons), and finally to the full sample across multiple timepoints. We compared model estimates for the commonly used *A*, *C*, and *E* components, as well as a subject-level component (*S*) in the longitudinal data, and the twin component (*T*), which captured the variance attributable to variance in the common environment of twin pairs. In addition, we explored the change in model estimates and model fit when adjusting for specific fixed effect covariates, and when excluding the twin sub-sample.

## Methods

### Sample

The ABCD Study^®^ is a longitudinal cohort of 11,880 adolescents beginning when participants were aged 9-11 years, with annual visits to assess mental and physical health (Volkow et al. 2018). The study sample spans 21 data acquisition sites and includes participants from demographically diverse backgrounds such that the sample demographics approximate the demographics of the United States (Garavan et al. 2018). The sample includes many siblings as well as a twin sub-sample consisting of 840 pairs of same-sex twins recruited from state birth registries at four sites (Garavan et al. 2018). Exclusion criteria for participation in the ABCD Study^®^ were: 1) lack of English proficiency in the child; 2) the presence of severe sensory, neurological, medical or intellectual limitations that would inhibit the child’s ability to comply with the study protocol; 3) an inability to complete an MRI scan at baseline. The study protocols were approved by the University of California, San Diego Institutional Review Board. Parent/caregiver permission and child assent were obtained from each participant. The data used in this study were obtained from ABCD Study^®^ data release 4.0.

Statistical analyses were conducted on a sample that included a total of 13,984 observations from 9,742 unique participants across two timepoints (the baseline and year 2 visits). The twin sub-sample used in this study consisted of 462 pairs of twins with complete data (258 DZ pairs, 204 MZ pairs; total *N* = 924). Observations were included in the final sample if the participant had complete data across sociodemographic factors (household income, highest parental education), available genetic data (to provide ancestry information using the top 10 principal components), and the phenotypes of interest. Table 1 shows the baseline demographics of the full sample as well as the twin sub-sample. Compared to the full sample, the twin sub-sample had a higher percentage of parents with bachelor’s degrees or above (67.3% compared to 61.8% in the full sample), and household income was shifted higher (52.7% with income over $100,000 compared to 42.0% in the full sample).

**Table 1.**
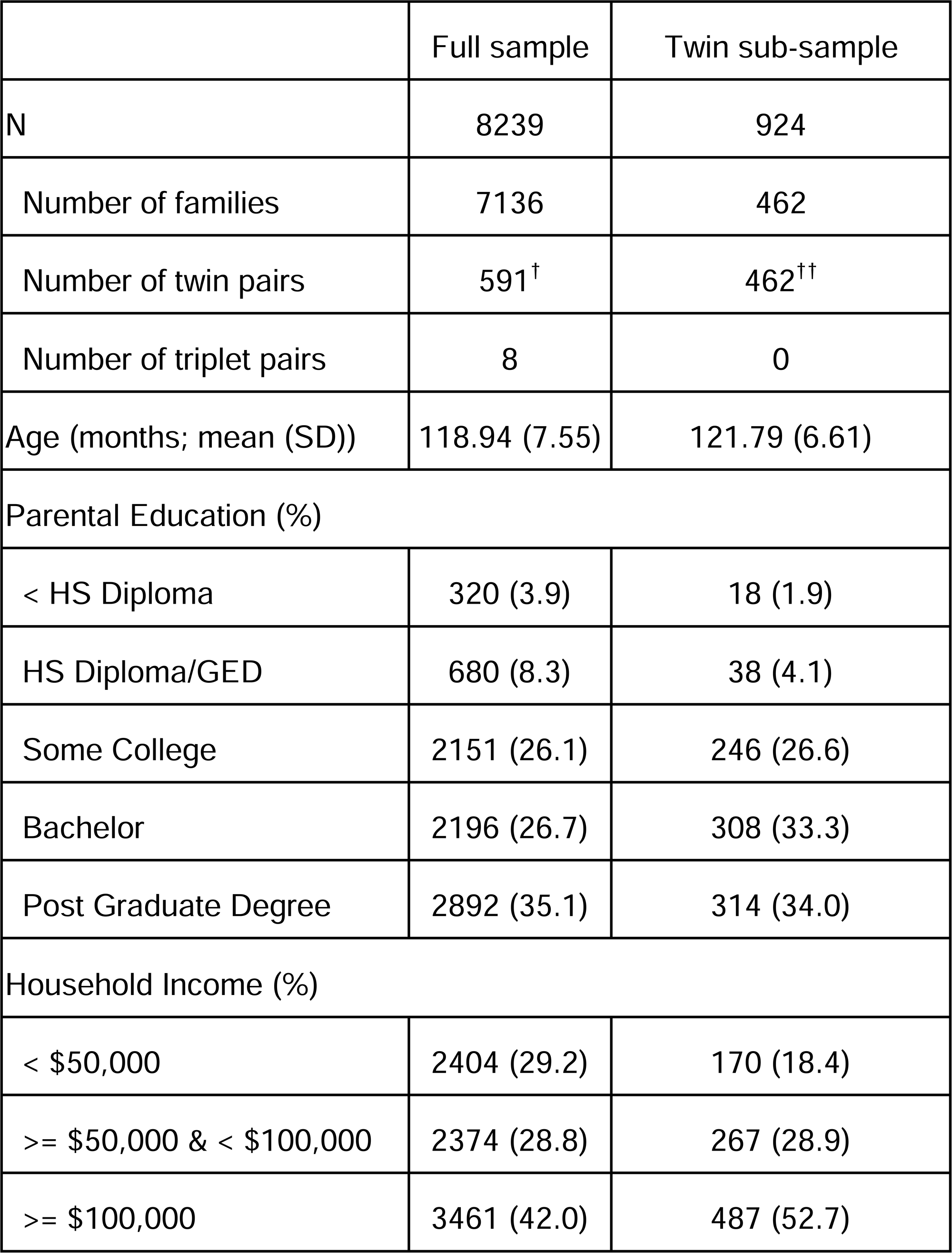
Sample information at baseline. All samples include complete cases only. ^†^Full sample included 241 pairs with SNP-derived relatedness > 0.9, implying 350 DZ pairs and 241 MZ pairs. ^††^Twin sub-sample included 258 DZ pairs and 204 MZ pairs.

## Measures

### Phenotypes of interest

For the present study, we included height as a phenotype of interest due to its common use in twin and family studies (Silventoinen et al. 2003), as well as the availability of larger genetic studies from samples of unrelated participants (Yengo et al. 2022). Several cognitive phenotypes were included from the NIH toolbox cognition battery (Gershon et al. 2013): specifically, we analyzed the raw composite scores measuring fluid and crystallized intelligence, which have been validated against gold-standard measures of cognition (Akshoomoff et al. 2013; Heaton et al. 2014). We also included the uncorrected scores from the flanker task, picture sequence memory task, list sorting memory task, pattern comparison processing speed, dimensional change card sort task (components of fluid cognition); and the oral reading recognition task and picture vocabulary task (components of crystallized cognition). In addition to the NIH Toolbox, we included the matrix reasoning test from the Wechsler Intelligence Scales for Children (WISC-V; Wechsler 2014), the total percent correct from the Little Man visuospatial processing task (Acker 1982), and the total number of items correctly recalled across the five learning trials of the Rey Auditory Verbal Learning Task (RAVLT; Schmidt 1996). See Extended Methods for a complete description of each phenotype of interest including data collection procedures.

### Covariates

Unless otherwise specified, models were run on data that was pre-residualized for age and sex only, in keeping with common practice for twin studies (Neale and Maes 2004). In models that included pre-residualization for additional covariates, these were chosen based on common practices in cognitive and behavioral research, and included recruitment site, parental education, household income, and the first ten genetic principal components.

### Genetic Principal Components and Genetic Relatedness

Methods for collecting genetic data have been described in detail elsewhere (Uban et al. 2018). Briefly, a saliva sample was collected at the baseline visit, as well as a blood sample from twin pairs. The Smokescreen™ Genotyping array (Baurley et al. 2016) was used to assay over 300,000 SNPs. Resulting genotyped and imputed SNPs were used for principal components derivation as well as genetic relatedness calculation.

The genetic principal components were calculated using PC-AiR (Conomos et al. 2015). PC-AiR was designed for robust population structure inference in the presence of known or cryptic relatedness. Briefly, PC-AiR captures ancestry information that is not confounded by relatedness by finding a set of unrelated individuals in the sample that have the highest divergent ancestry and computes the PCs in this set; the remaining related individuals are then projected into this space. This method has been recommended by the Population Architecture through Genomics and Environment Consortium (Wojcik et al. 2019), which is principally concerned with conducting genetic studies in diverse ancestry populations.

PC-AiR was run on using the default suggested parameters from the GENESIS package (Gogarten et al. 2019). We used non-imputed SNPs passing quality control (516,598 variants and 11,389 individuals). Using the computed kinship matrix, PC-Air was then run on a pruned set of 158,103 SNPs, which resulted in 8,005 unrelated individuals from which PCs were derived – leaving 3,384 related individuals being projected onto this space.

We then computed a GRM using PC-Relate (Conomos et al. 2016). PC-Relate aims to compute a GRM that is independent from ancestry effects as derived from PC-AiR. PC-Relate was run on the same pruned set of SNPs described above using the first two PCs computed from PC-Air.

## Data analysis

### Pre-residualization

We used R version 3.6.3 for data processing. After obtaining the sample of complete cases for all variables, phenotypes were pre-residualized for age and sex using the *lm* function. For certain models (see Table 2), we additionally included the following covariates during this residualization step: site, parental education, income, and the first ten genetic principal components. The purpose of pre-residualization was to ensure that both FEMA and *OpenMx* implementations were fitting random effects to the same data. Because our models only fit random effects, and because FEMA implements an unbiased estimation of total variance, the FEMA implementation was therefore mathematically equivalent to *OpenMx*.

**Table 2.**
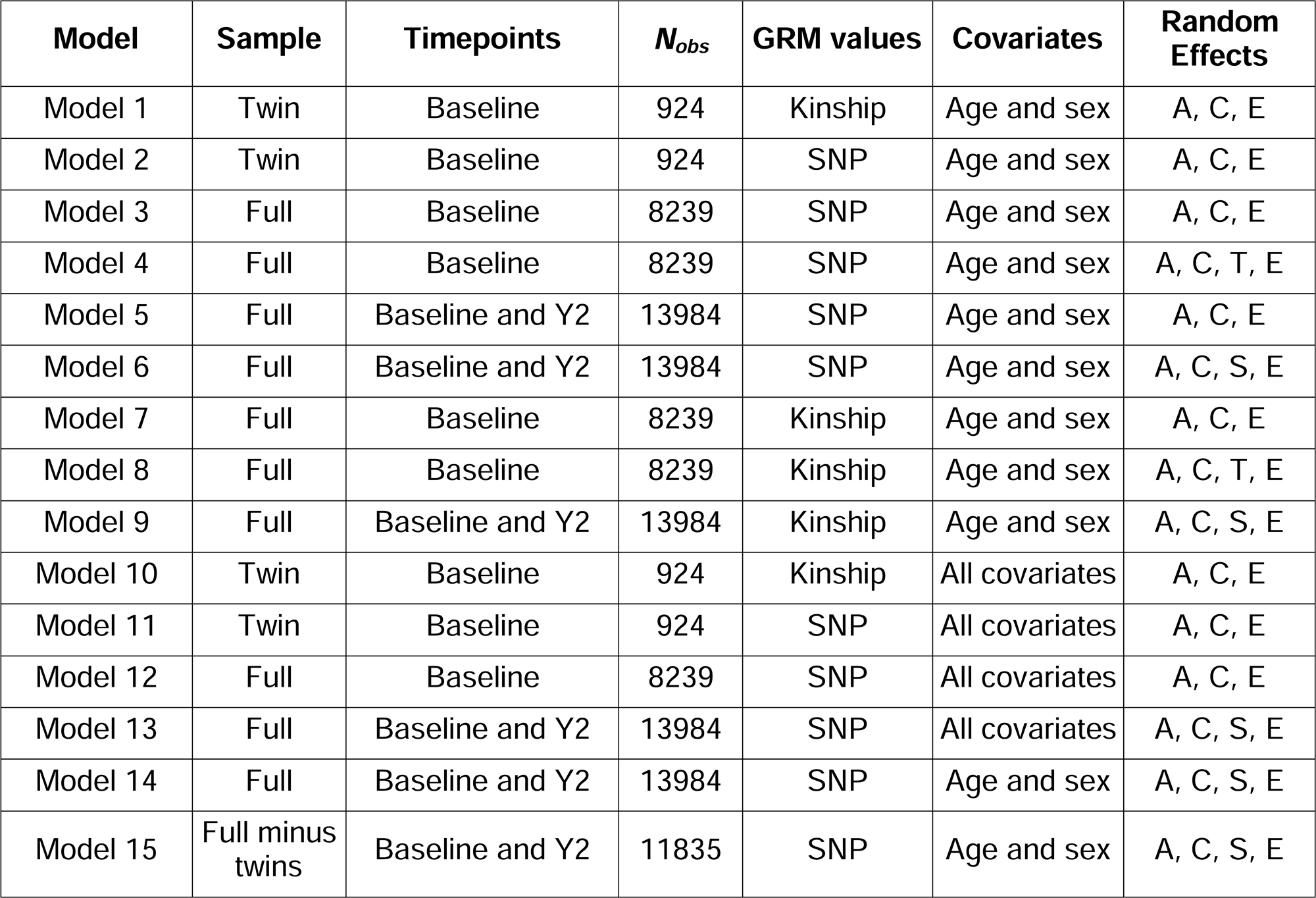
List of model specifications. GRM = genetic relatedness matrix (derived from SNP data or kinship data); N_obs_ = number of observations; SNP = single nucleotide polymorphism; Y2 = year 2 follow-up visit. All models pre-residualized for age and sex; when specified, “all covariates” includes these as well as site, parental education, household income, and first ten genetic principal components. Samples correspond to sample information listed in Table 1. Random effects: A = additive genetic relatedness, C = common environment, S = subject, T = twin status (shared pregnancy ID), E = unexplained variance / error.

Previous work has found evidence for a practice effect in some of the cognitive measures from the ABCD Study^®^ (Anokhin et al. 2022). Therefore, in models that included data from baseline and year 2, we included a “practice effect” as a dummy variable in the pre-residualization step. This variable was equal to 0 if the observation was the first instance of data for that participant (i.e., all participants had 0 at baseline), and 1 if the participants were providing data for the second time at the year 2 visit. Most participants (*N* = 4242, 76.19%) had a value of 1 at the year 2 visit.

### Model specification

We ran a series of models, described in Table 2. For each model, we specified whether genetic relatedness was “SNP-derived” (calculated using PC-AiR and PC-Relate; Conomos et al. 2015; Conomos et al. 2016) or “kinship-derived”. For kinship-derived relatedness, we used the zygosity data from the twin sub-sample to assign a value of 1 for MZ twins, 0.5 for DZ twins, and 0.5 for all other siblings (under the assumption that there are only full siblings within a family).

Since each phenotype was pre-residualized, we only needed to estimate the random effects components in each LME run within FEMA and *OpenMx*. These included an effect of family ID (common environment, *C*), additive effect of genetic relatedness (*A*), subject (*S*), twin status (*T*, calculated by creating a variable “pregnancy ID” that was shared by any two individuals with the same family ID and same birth date), and unique environment/unexplained variance (*E*).

### OpenMx

We first ran an ACE model in the baseline twin sample, using the *OpenMx* package in R (package version 2.20.6; Neale et al. 2016). We elected to use *OpenMx* as the comparison software due to its widespread use in twin and family studies to estimate heritability. We chose to use the restricted maximum likelihood (REML) estimator within *OpenMx*, which differs from ML estimators by a) using an unbiased estimation to calculate total variance, and b) first estimating the random effects iteratively and then estimating the fixed effects coefficients, as opposed to alternating estimation of variances and fixed effects. However, due to the preresidualization step described above, in our models we solely estimated random effects, such that the REML estimator in *OpenMx* provided a good comparison for FEMA (a ML estimator that uses an unbiased estimation of total variance). We ran *OpenMx* using R version 3.6.3., using the default SLSQP optimizer. Because data were preresidualized for age and sex, we did not fit any additional covariates. *OpenMx* provides likelihood-based confidence intervals by default (Neale and Miller 1997) which we used to compare with the likelihood-based confidence intervals calculated in FEMA.

### Fast Efficient Mixed Effects Analysis (FEMA)

FEMA was developed for the efficient implementation of mass univariate LMEs in high dimensional data (e.g., brain imaging phenotypes; Fan et al. 2021). Whereas the original version of FEMA used a method of moments estimator for increased computational efficiency, we modified the package to allow the user to select a ML estimator. When users specify ML as the random estimator, FEMA arrives at parameter estimates by minimizing the log likelihood function specified in *FEMA_loglik*. An updated version of FEMA, including the relevant code, is available at the time of this publication (https://github.com/cmig-research-group/cmig_tools). Because FEMA uses an unbiased estimation of total variance, and we were only fitting random effects and not fixed effects, the estimates from the FEMA implementation of ML regression were predicted to be mathematically equivalent to the REML estimator used in *OpenMx*. To run FEMA, we passed a design matrix (the design matrix was “empty” because we were not fitting any fixed effects) as well as a file containing a matrix of (SNP- or kinship-derived) genetic relatedness values. FEMA then used a nested random effects design to create a sparse relatedness matrix, in which the relatedness values for all participants not assigned the same family ID was set to zero. For all models we reported the random effects variances as a percent of the total variance in the residualized phenotype that was explained by variance in the random effect of interest. As a result, for a given model, the variance component estimates sum to 1 (representing 100% of the variance in the residualized phenotype). For ease of interpretation and comparison to previous literature, in this paper the term “heritability estimate” refers to the percent of residualized phenotypic variance that is explained by variance in genetic relatedness, i.e., variance explained by variance in *A*.

### Confidence Interval Calculation

To generate 95% confidence intervals around parameter estimates, we used the *mxCI* function within *OpenMx*. The likelihood-based confidence intervals returned using *mxCI* are obtained by increasing or decreasing the value of each parameter until the −2 log likelihood of the model increases by an amount corresponding to the requested interval. The implementation of likelihood-based confidence intervals has been described in detail by Neale and Miller (1997).

FEMA used the same profile likelihood method to calculate confidence intervals (see Sprott 2000). Code for calculating confidence intervals is available within the *FEMA_fit* function on the publicly available GitHub repository. Briefly, *FEMA_fit* calculates a log likelihood threshold corresponding to the requested confidence interval, then solves a quadratic equation that describes the change in likelihood as a function of change in parameter value. The solution to this equation is applied to the parameter value of interest to achieve the specified confidence interval. Differences between FEMA and *OpenMx* confidence interval calculation are likely to be due to small differences in the specific implementations used.

### Model Comparison

For comparing two models that used identical samples, we calculated the Akaike Information Criterion (AIC) as (Akaike 1974):

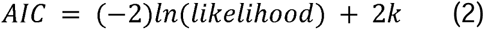

where *k* represents the number of model parameters. Therefore, the difference in AIC between two models (Δ*AIC*) can be calculated as:

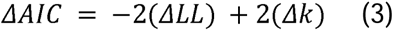

where Δ*LL* represents the difference in log likelihood between the two models and Δ*k* represents the difference in the number of parameters between the two models. In models that have the same level of complexity (Δ*k* = 0), the Δ*AIC* is equal to −2(Δ*LL*). We chose to use the AIC as opposed to the likelihood ratio test statistic for model comparison because several comparisons were not between nested models.

## Results

### ACE model (Model 1) in FEMA versus *OpenMx*

A summary of heritability estimates (i.e., the *A* random effects) from all models is provided in Supplementary Table 1. To compare model estimates between *OpenMx* and FEMA, we fit the same ACE model in each, using the same sample of 462 complete twin pairs from the twin sub-sample (i.e., pairs in which each twin had complete data for all phenotypes). Figure 1 shows a comparison of the two results as well as the parameter estimates using FEMA. The difference in heritability estimates between the two software packages was less than 0.001 for all phenotypes. On comparing these models (Figure 1B), we found that the difference in AIC was less than 0.05 for all phenotypes, indicating that there was no difference in the model fit. Because the model estimates and model fits were practically the same, we concluded that the two implementations were indeed mathematically equivalent, and therefore elected to use the LME implementation in FEMA for all further analyses.

**Figure 1.**
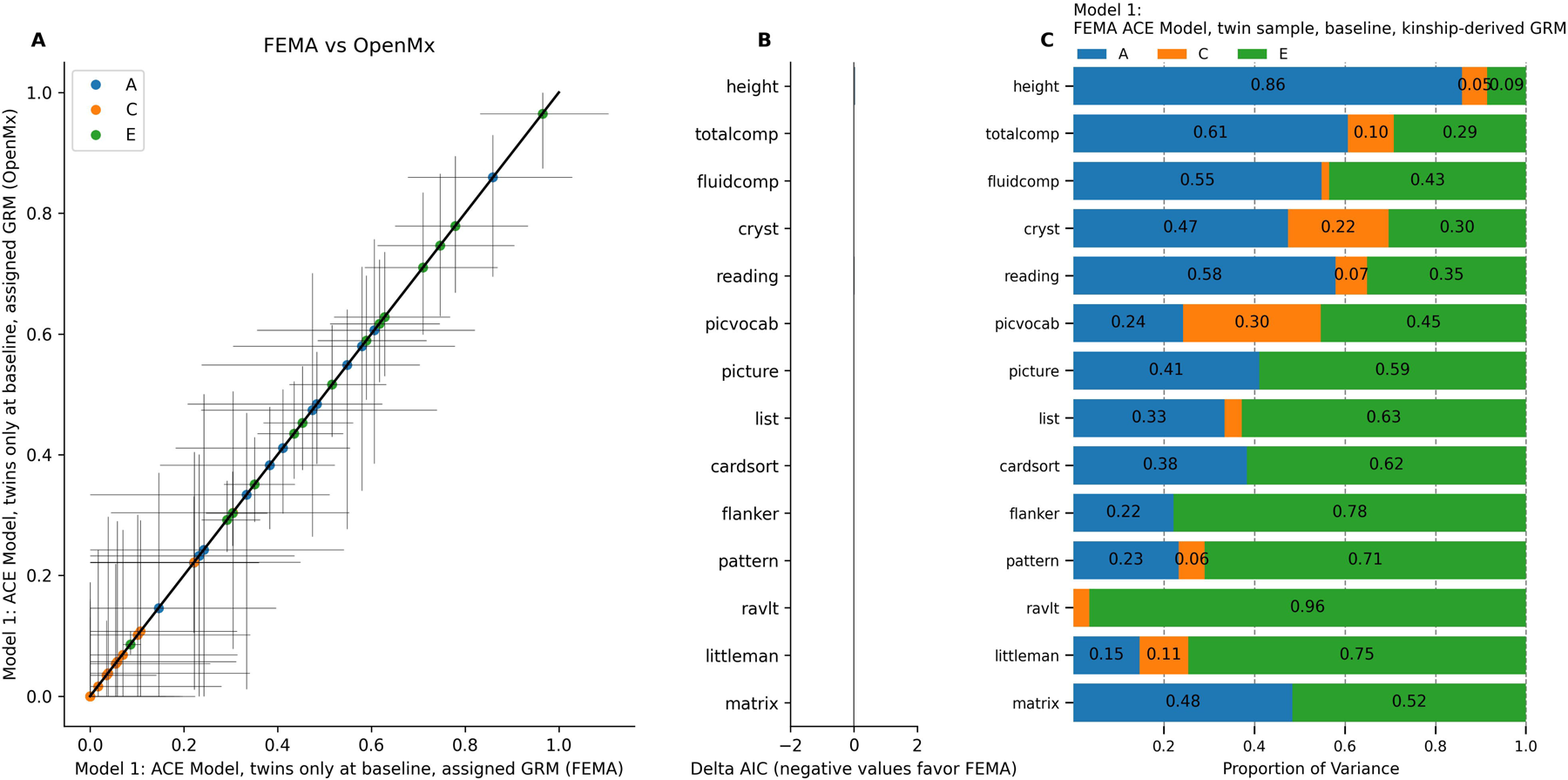
ACE Model (Model 1) in FEMA versus OpenMx, twin sub-sample at baseline. A) Comparison of model estimates. Horizontal error bars represent confidence interval calculated in FEMA; vertical error bars represent confidence intervals calculated in OpenMx. B) Difference in Akaike Information Criterion in FEMA versus in OpenMx. C) Random effects estimates from FEMA.

### Effect of including measured genetic relatedness (Model 2)

To test whether variance component estimates differ when including SNP-versus kinship-derived genetic relatedness, we fit two versions of the ACE model in the baseline twin sample. Model 1 (the ACE model described above, implemented in FEMA) used a matrix of kinship-derived relatedness values (1.0 for MZ twins and 0.5 for DZ twins) whereas Model 2 used a matrix of SNP-derived relatedness values.

The models provided equivalent heritability estimates, with differences in A estimates ranging from −0.01 (Little Man Task) to 0.03 (pattern comparison; Figure 2A). On inspecting the differences in the AIC between the two models, we found that using SNP-derived GRM led to small improvements in the overall model fit. This improvement was most pronounced for height (Δ*AIC* = −1.04) but less so for the cognitive phenotypes (Figure 2B). Random effects variance component estimates are presented in Figure 2C; overall, using SNP-derived GRM did not lead to dramatically different parameter estimates than using kinship-derived GRM.

**Figure 2.**
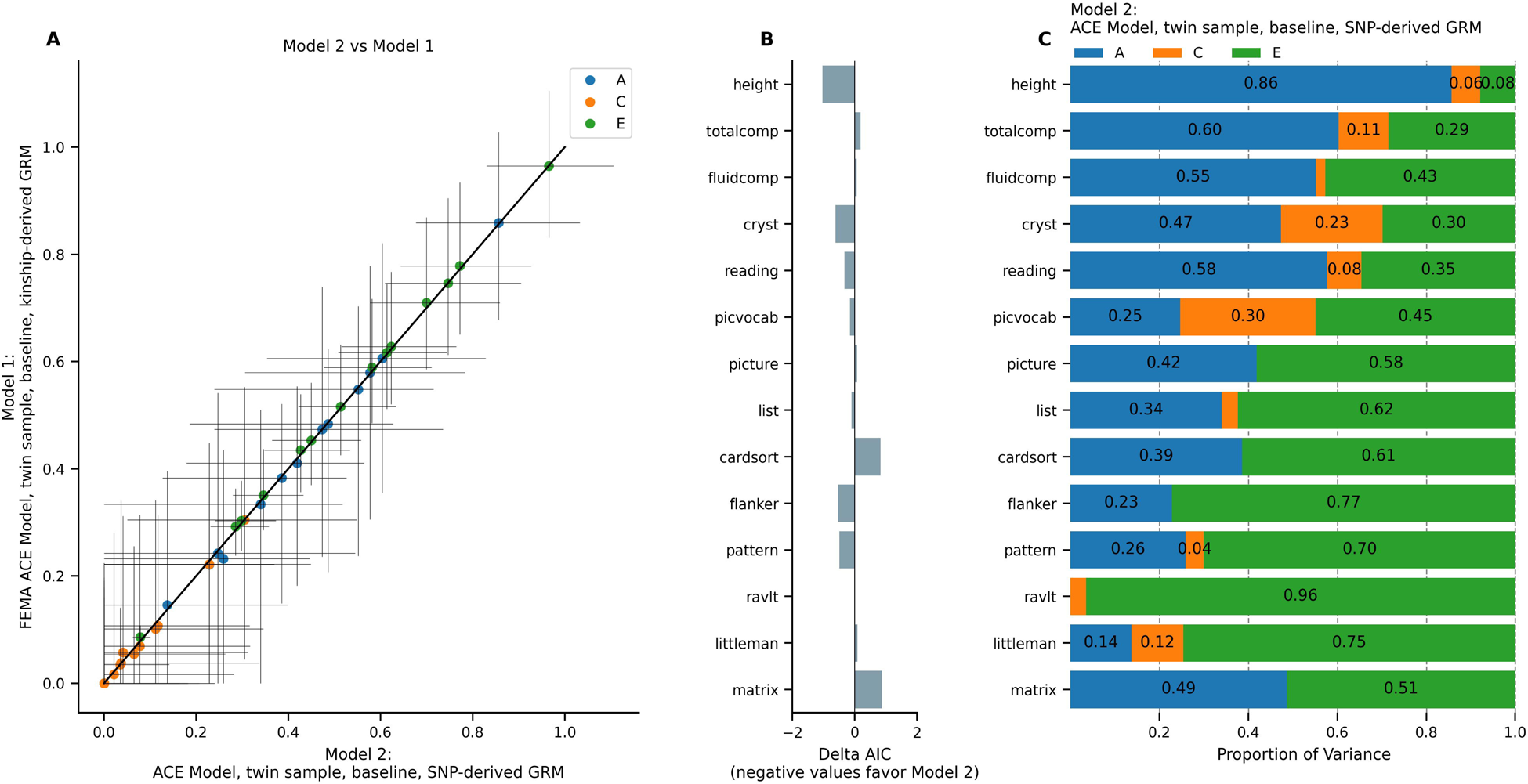
ACE Model using kinship-derived (Model 1) versus SNP-derived GRM (Model 2), twin sub-sample at baseline. A) Comparison of model estimates. Horizontal error bars represent confidence interval calculated in Model 2; vertical error bars represent confidence intervals calculated in Model 1. B) Difference in Akaike Information Criterion in Model 2 versus in Model 1. C) Random effects estimates from Model 2.

### Effect of increased sample size (Model 3)

To examine how the variance component estimates in the twin sample (*n* = 924) differ from the full ABCD Study^®^ baseline sample (*n* = 8,239), we next compared the ACE model in these two groups. Model 2 (from previous analysis) and Model 3 both used the SNP-derived GRM values and included *A*, *C*, and *E* random effects. The two models are therefore equivalent except for the much larger sample fit in Model 3. As described in Methods, the sparse clustering method within FEMA ignored the genetic relatedness among individuals with different family IDs. In practice, this meant that the sample of 8,239 unique subjects at baseline was clustered into 7,136 families, and genetic relatedness values were only used for individuals within families.

Figure 3 shows the estimates from Model 3 and their comparison to Model 2. The increased sample size led to smaller confidence intervals for most random effects estimates calculated in Model 3 (Figure 3A). The changes in heritability estimates ranged from −0.31 (NIH Toolbox Fluid Cognition) to +0.04 (height). The estimated total variance was larger in Model 3 for most phenotypes (Figure 3B), with the largest increase in variance in total composite cognition (24.36% increase in total variance), crystallized cognition (23.76% increase), and oral reading recognition (24.08% increase). Because the two models were fit to different samples, it was not possible to directly compare their AIC model fit from the likelihood statistics. Figure 3C shows the random effects variances from Model 3; results indicate that the total variance in the full sample was larger than that in the twin sample, and using the full sample led to smaller estimates of A and larger estimates of C compared to the twin sample.

**Figure 3.**
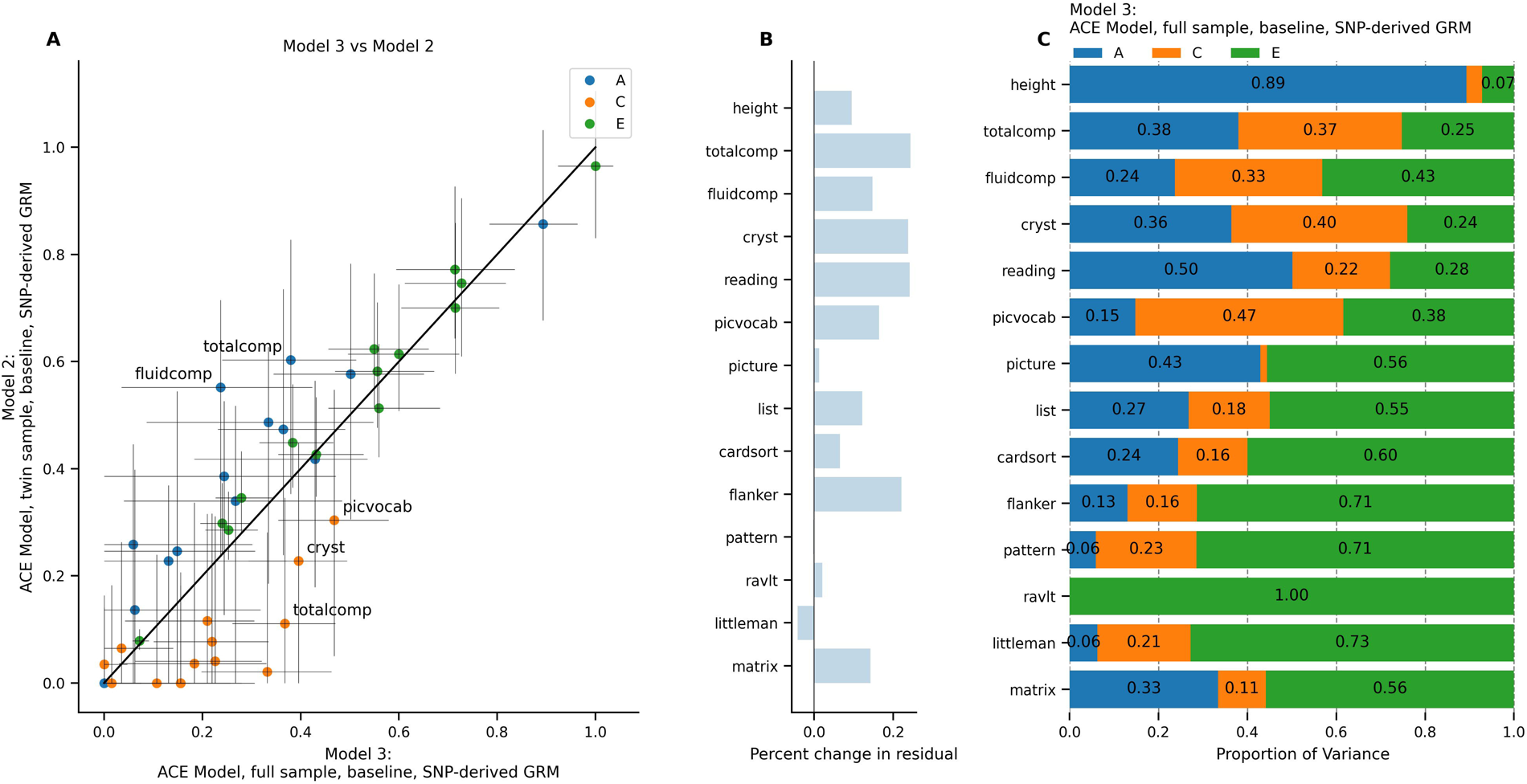
ACE Model using full sample (Model 3) compared to twin sub-sample (Model 2) at baseline. A) Comparison of model estimates. Horizontal error bars represent confidence interval calculated in Model 3; vertical error bars represent confidence intervals calculated in Model 2. B) Difference in total residual variance in Model 3 versus in Model 2. C) Random effects estimates from Model 3.

### Adding a Twin random effect (Model 4)

Given that the full sample analysis included singletons, half siblings, and adopted siblings, as well as twins and triplets, we next tested whether the addition of a random effect of twin status (*T*) led to a change in parameter estimates. We calculated a “pregnancy ID” that was shared by individuals who had the same family ID and the same birth date. We then used this “pregnancy ID” to code for the *T* random effect in an ACTE Model (Model 4). Supplementary Figure 1 shows the model estimates from Model 4 as well as a comparison to Model 3; the two models are equivalent with the exception of the *T* random effect.

For most phenotypes, the addition of the *T* random effect did not lead to a change in parameter estimates (i.e., *T* was estimated to be 0). The largest change in parameter estimates was in matrix reasoning (heritability estimate decreased by 0.05, *T* estimated at 0.05; Supplementary Figure 1A). Model comparison found that the difference in the AIC was at or near 2.0 for all phenotypes except for the RAVLT (Δ*AIC* = 1.68) and matrix reasoning (Δ*AIC* = 1.35). Because the AIC was calculated as *-2*Δ*LL* plus double the difference in model parameters (Equation 3), the consistent values of 2.0 reflect that the *-2*Δ*LL* statistic was approximately 0 before the penalization for the additional parameter in Model 4 (Supplementary Figure 1B). The random effects variances for Model 4 are shown in Supplementary Figure 1C. Results indicate that T did not explain residual variance above and beyond the inclusion of A and C (matrix reasoning being a notable exception to this pattern).

### Incorporation of two timepoints (Model 5, 6)

To compare the difference between cross-sectional and longitudinal samples, we next moved from examining the full sample at baseline (Model 3) to the full sample at baseline and Year 2 (Model 5). Models 3 and 5 were equivalent except for the difference in sample size (i.e., Model 5 did not account for nesting of data within subjects, in order to directly assess this effect in Model 6). Because not all phenotypes were available at the Year 2 visit, models that included baseline and Year 2 data only included pattern comparison processing speed, flanker task performance, picture sequence memory, picture vocabulary, oral reading recognition, crystallized cognition, RAVLT, Little Man Task, and height. To account for nesting of multiple visits within subjects, we added a random effect of subject (*S*) in Model 6. Figure 4 shows the change in random effects variances moving from Model 3 to Model 5 and from Model 5 to Model 6.

**Figure 4.**
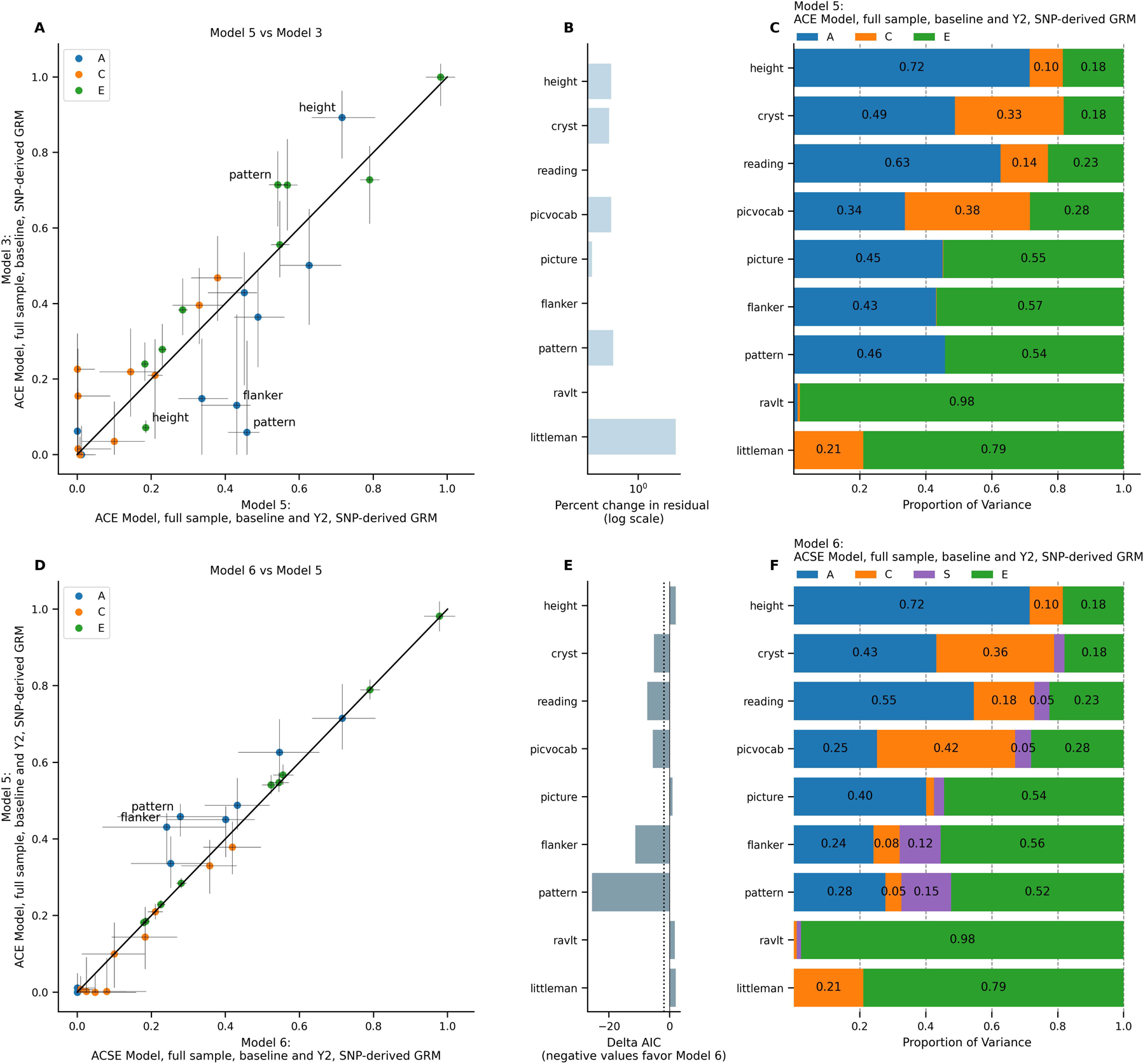
ACE and ACSE Model in full sample, baseline only (Model 3) versus baseline + year 2 (Models 5 and 6). A) Comparison of estimates from Model 5 versus Model 3. Horizontal error bars represent confidence interval calculated in Model 5; vertical error bars represent confidence intervals calculated in Model 3. B) Difference in total residual variance in Model 5 versus in Model 3. C) Random effects estimates from Model 5. D) Comparison of estimates from Model 6 versus Model 5. Horizontal error bars represent confidence interval calculated in Model 6; vertical error bars represent confidence intervals calculated in Model 5. E) Difference in Akaike Information Criterion in Model 6 versus in Model 5. F) Random effects estimates from Model 6.

Adding the second visit led to overall increases in the heritability estimates for the cognitive phenotypes, with changes ranging from −0.06 (Little Man Task) to +0.40 (pattern comparison; Figure 4A). Conversely, the heritability estimate for height decreased by 0.18. Expanding from the full sample at baseline to the full sample at baseline and Year 2 led to an increase in the total residual variance for several phenotypes; the most notable increase was in the Little Man Task (7928% increase in total residual variance). Adding the random effect of subject led to minimal change (<0.02) in the heritability estimate for height, Little Man Task, and RAVLT, but a decrease in heritability estimates across the other cognitive phenotypes (changes ranging from −0.19 to −0.05) compared to estimates from Model 5. The total difference in heritability estimates going from Model 3 to Model 6 ranged from −0.18 (height) to +0.22 (pattern comparison; Figure 4D).

Model comparison between Model 5 and Model 6 found that the model was substantially improved for crystallized cognition (Δ*AIC* = −5.19), oral reading recognition (Δ*AIC* = −7.45), picture vocabulary (Δ*AIC* = −5.61), flanker (Δ*AIC* = −11.33), and pattern comparison (Δ*AIC* = −25.60). However, the difference in the fit was smaller for height (Δ*AIC* = +2.00), picture sequence memory (Δ*AIC* = +0.89), the RAVLT (Δ*AIC* =+1.72), and the Little Man Task (Δ*AIC* =+2.00; Figure 4E). Figure 4C and 4F show the random effects variances from Model 5 and Model 6. Overall, the increased sample size led to increased total residual variance and narrower confidence intervals, increases in heritability estimates for cognitive phenotypes, and a decrease in the heritability estimate for height, compared to the baseline only sample.

### Effect of using kinship-derived genetic relatedness in large samples (Model 7-9)

For the next set of models, we aimed to approximate a cohort study design in which genetic data were not available, to examine whether model estimates using several thousand subjects (Models 3,4,6) changed in the absence of SNP data. We used a matrix of kinship-derived genetic relatedness (assigning 1.0 for MZ twins from the twin sub-sample, and 0.5 for DZ twins from the twin sub-sample and all other individuals in the same family). The kinship-derived relatedness value therefore assumed that all non-twins in the same family, as well as twins who were not part of the twin sub-sample, were full siblings.

Figure 5 compares the ACSE longitudinal model with an equivalent model that used kinship-derived genetic relatedness. Supplementary Figure 2 shows the same question of kinship-derived versus SNP-derived relatedness applied to Models 3 and 4. Overall, the random effects estimates were largely unchanged with the use of kinship-derived GRM, with the largest changes in the ACSE model occurring in flanker (Δ*A* = 0.09) and pattern comparison (Δ*A* = −0.07; Figure 5A, Supplementary Figure 2A,D). Model comparison using Δ*AIC* found that the ACSE model using SNP-derived GRM had better model fit for height (Δ*AIC* = −36.36), crystallized cognition (Δ*AIC* = −17.59), oral reading recognition (Δ*AIC* = −19.45), picture vocabulary (Δ*AIC* = −8.75), and pattern comparison (Δ*AIC* = −8.84) compared to the model using kinship-derived GRM; the difference in model fit was less pronounced for picture sequence memory (Δ*AIC* = +0.46) and flanker (Δ*AIC* = +0.45; Figure 5B, Supplementary Figure 2B,E). Figure 5C and Supplementary Figure 2C and 2F show the random effects variances for models using kinship-derived genetic relatedness; results indicate that using SNP-derived versus kinship-derived GRM led to small changes in heritability estimates, though model fit was better with SNP-derived GRM.

**Figure 5.**
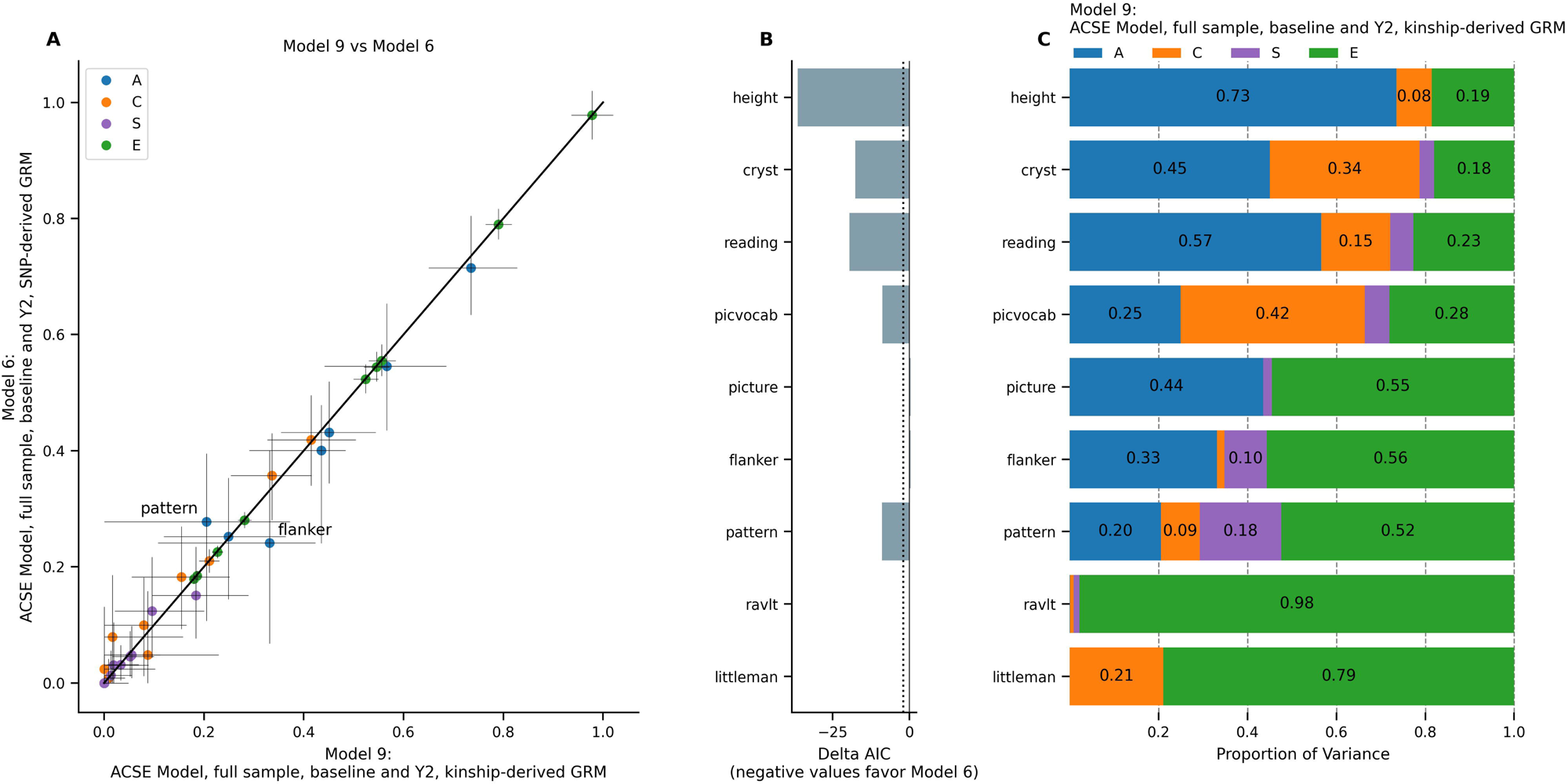
ACSE model using SNP-derived (Model 6) versus kinship-derived genetic relatedness (Model 9), full sample, baseline + year 2. A) Comparison of estimates from Model 9 versus Model 6. Horizontal error bars represent confidence interval calculated in Model 9; vertical error bars represent confidence intervals calculated in Model 6. B) Difference in Akaike Information Criterion in Model 9 versus in Model 6. C) Random effects estimates from Model 9.

### Residualizing for additional covariates (Models 10-13)

While it is common in twin and family analyses to include only age and sex as fixed effects, behavioral scientists often include additional fixed effects such as sociodemographic variables or recruitment site as covariates. To test whether the inclusion of such variables led to changes in our random effects estimates, we ran several of our original models with additional variables included in the pre-residualization step (i.e., site, parental education, income, and the first ten genetic principal components). Figure 6 shows the results of this model comparison applied to Model 1 (the “classic” ACE model). Supplementary Figure 3 shows the same pre-residualization and model comparison applied to Models 2-4 and 6. In the classic ACE model, the *A* estimate tended to decrease and the *C* estimate tended to decrease in the models that included additional covariates (Figure 6A). Residualizing for additional covariates led to a decrease in the total residual variance across all phenotypes, with decreases ranging from −2.67% (RAVLT) to −26.02% in the ACE model (crystallized cognition; Figure 6B). Because the two models were run on different datasets (pre-residualized for different covariates), we did not calculate the difference in AIC between the two models. Figure 6C and Supplementary Figure 3C, 3F, and 3I show the random effects variances for the models that were residualized for additional covariates. In general, the inclusion of additional fixed effects led to lower residual variance being attributed to the random effects of interest, and a lower proportion of the remaining variance was attributed to *A* and *C*.

**Figure 6.**
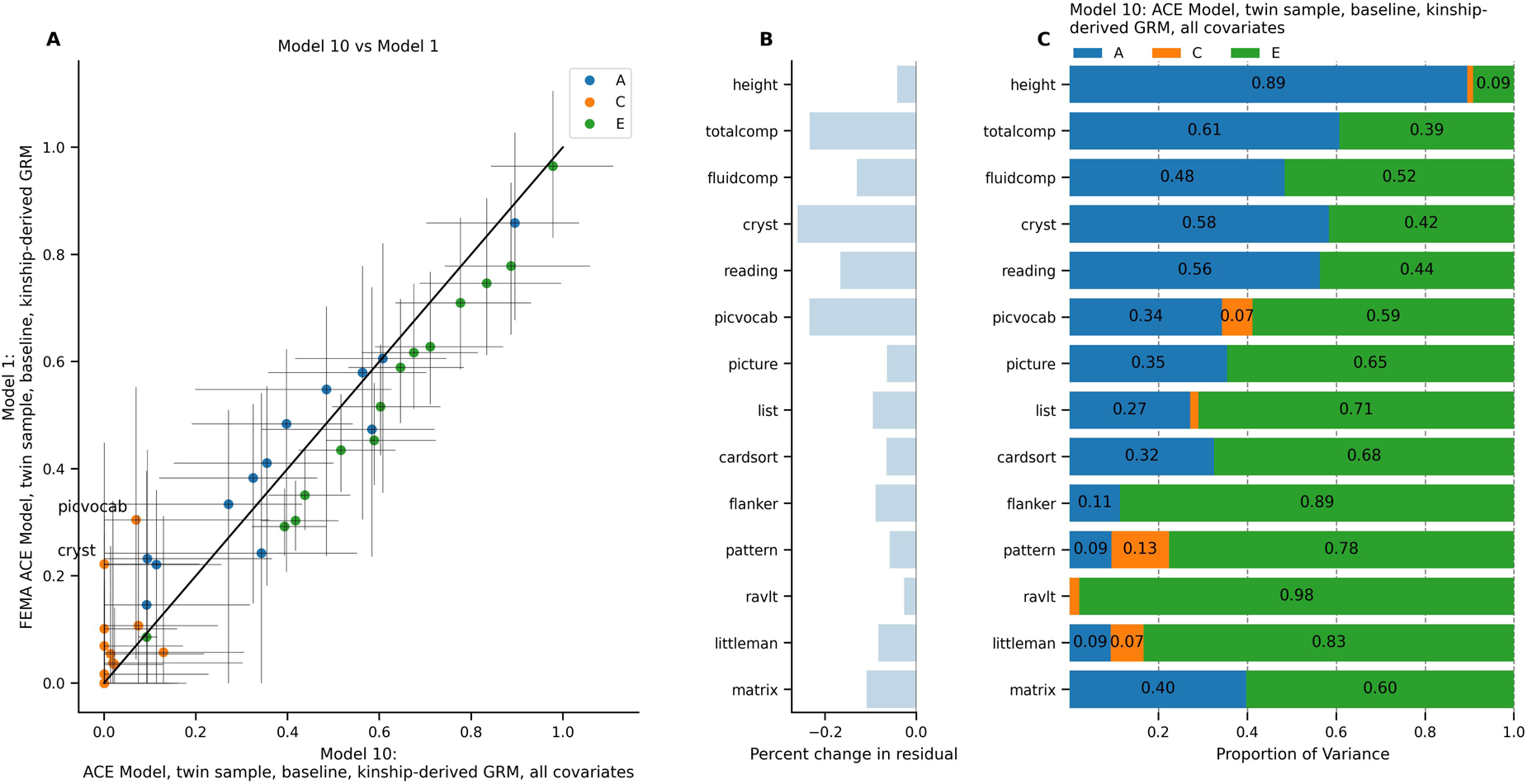
ACE model in twin sub-sample at baseline, residualizing for all covariates (Model 10) versus age and sex only (Model 1). A) Comparison of estimates from Model 10 versus Model 1. Horizontal error bars represent confidence interval calculated in Model 10; vertical error bars represent confidence intervals calculated in Model 1. B) Difference in total residual variance in Model 10 versus in Model 1. C) Random effects estimates from Model 10.

### Effect of removing the twin-enriched sample (Models 14-15)

The size and structure of the ABCD Study^®^ cohort, with its embedded twin sub-sample as well as the large number of related participants, led us to test the degree to which the model fit depended on having a large subset of MZ and DZ twins. This question was motivated by the fact that many large cohort studies do not include a specifically twin-enriched sample; we aimed to explore whether such studies can realistically perform similar models to those tested in this paper.

As a proxy for the general population, we removed the twin sub-sample. This left a small number of twins and triplets recruited through the general recruitment pipeline (168 twin pairs and 6 sets of triplets, with 57 pairs of participants with genetic relatedness > 0.9 across the full sample). The number of twin and triplet sets in this sample (174 out of 8131 pregnancies, 2.14%) was less than the 3.11% twin birth rate reported in the general population of the United States (Osterman et al. 2021). We therefore assumed that the ABCD Study^®^ sample excluding the embedded twin sub-sample was a proxy for a population sample with a naturally occurring number of twins. We then fit an ACSE model, applied to the full sample excluding the twin sub-sample, at baseline and year 2, to represent the “best” model possible of those explored thus far, excluding the *T* random effect (Model 15). We compared this model to the same ACSE model applied to the full sample, inclusive of twins (Model 14).

A comparison of the parameter estimates is shown in Figure 7A. The model excluding the twin sub-sample led to a difference in *A* estimates of −0.19 (picture sequence memory task) to +0.13 (pattern comparison). Excluding the twin sub-sample led to an increase in the total residual variance across all phenotypes, with changes ranging from +0.24% (pattern comparison) to +17.55% (Little Man Task; Figure 7B). Because the two models were fit to different samples, it was not possible to directly compare model fit from the likelihood statistics. Figure 7C shows the random effects variances from the model that omitted the twin sub-sample participants. Overall, excluding the twin sub-sample led to larger total residual variance, with varied effects on heritability estimates.

**Figure 7.**
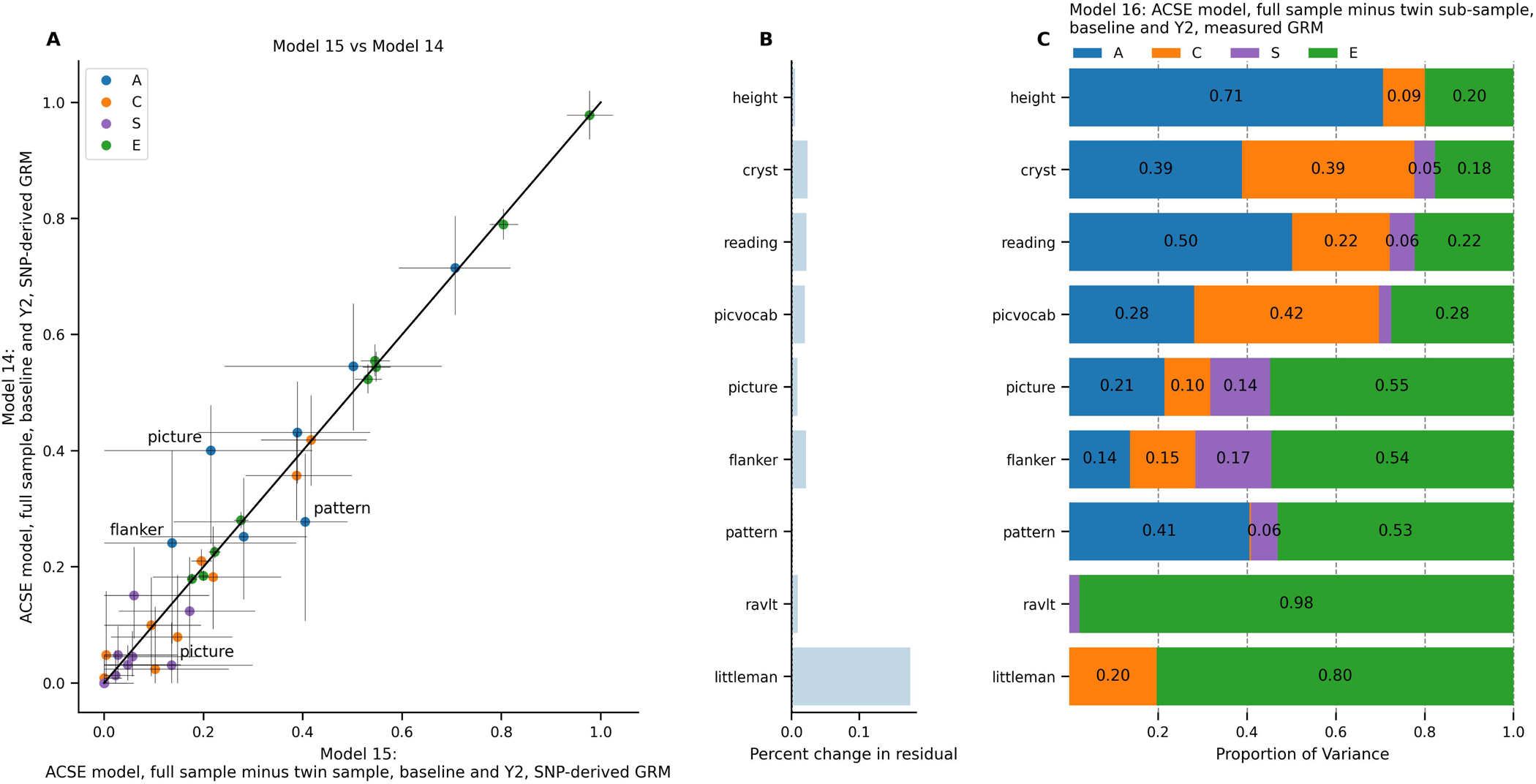
ACSE model in full sample omitting twin registry participants, baseline + year 2 longitudinal sample (Model 15). Comparison model is equivalent but includes the full sample inclusive of twin registry participants (Model 14). A) Comparison of estimates from Model 15 versus Model 14. Horizontal error bars represent confidence interval calculated in Model 15; vertical error bars represent confidence intervals calculated in Model 14. B) Difference in total residual variance in Model 15 versus in Model 14. C) Random effects estimates from Model 15.

## Discussion

In this paper we present results from different modeling strategies for implementing the ACE model using LMEs, as implemented in FEMA. FEMA is capable of applying the ACE model as well as incorporating additional features such as using a sparse matrix of within-family genetic relatedness and a random effect of subject to model longitudinal data. Notably, the use of FEMA to incorporate relatedness across all subjects within a family allows for the flexibility to include the full ABCD Study^®^ sample, rather than restricting analysis to the twin sub-sample. Future updates to the ABCD Study^®^ Data Exploration and Analysis Portal will incorporate an online implementation of FEMA so that users can specify mixed effects models of interest using the ABCD Study^®^ dataset. In addition, future analyses will incorporate random effects estimates across the high-dimensional brain imaging data present in the ABCD Study^®^.

We first applied the ACE model in the baseline twin sample. FEMA and *OpenMx* found nearly equivalent estimates for all random effects variances, demonstrating the equivalence of the two models being fitted. We estimated the heritability of height at 0.86, which is near the top of the range of twin heritability estimates reported by a comparative study of twin cohorts in eight countries (ranging from 0.68 to 0.87; Silventoinen et al. 2003). Our twin heritability estimate for height was higher than the SNP heritability, which was recently estimated to be 40% of phenotypic variance in European ancestry populations and 10%-20% in other ancestries (Yengo et al. 2022). Of the cognitive phenotypes, we found the highest twin heritability estimate for total composite cognition (0.61) and oral reading recognition (0.58), consistent with prior findings that heritability estimates tend to be higher for more “crystallized” and culturally sensitive measures of cognition (Kan et al. 2013). Interestingly, the picture vocabulary test had a relatively lower heritability estimate in this model (0.24) compared to the reading recognition test (0.58), which may reflect a difference in the cultural sensitivity of the two “crystallized” cognition tasks. The NIH Toolbox tasks comprising fluid cognition (flanker task, picture sequence memory task, list sorting, pattern comparison, and dimensional card sort) ranged in heritability estimates from 0.22 (flanker) to 0.41 (picture), which is within the wide range of heritability estimates for similar tasks in children (approximately 0-0.6; see Kan et al. 2013). Interestingly, the RAVLT had near-zero estimates for all random effects variances in all models, indicating that this task may be exceptionally unreliable in this sample, or perhaps particularly prone to variance in measurement. For further information regarding the heritability of 14,500 phenotypes in the ABCD Study^®^ baseline sample, see recent work by Maes and colleagues (2023).

We next tested the change in model fit and parameter estimation when using SNP-derived genetic relatedness rather than kinship-derived relatedness. Parameter estimates were largely unchanged, reflecting that in a twin sample, the kinship-derived relatedness values of 0.5 and 1 are sufficient to arrive at similar random effects estimates compared to models using SNP-derived relatedness (though the model fit was improved with the SNP-derived relatedness values for several phenotypes). Based on these results, researchers who are deciding between using SNP-derived or kinship-derived relatedness data to model cognitive phenotypes may prefer to choose based on practical considerations (such as availability of different data types, or the relevance of each relatedness estimate to the research question of interest) rather than making a categorical decision based on model fit.

Perhaps one of the most exciting applications comes when extending the model to the full ABCD Study^®^ sample. By leveraging the sparse clustering method used by FEMA to handle genetic relatedness only for participants within families, we were able to take advantage of the diverse distribution of genetic relatedness, ranging from 0 (e.g. adopted siblings) to 1 (i.e., MZ twins) for any pair of participants within a family. Unlike the large computational load generated by other similar genome-based REML regressions, the use of sparse clusters allowed FEMA to dramatically cut the computational time (Fan et al. 2021), allowing all the analyses in this paper to be fit on a single machine without the use of parallel computing. Using the full sample, first at baseline then with the addition of the Year 2 data, led to narrower confidence intervals, as shown in Figure 3. Inclusion of the full sample led to lower heritability estimates for several cognitive phenotypes, which may be related to the relative homogeneity of the twin sub-sample leading to potential for overestimation of heritability. Of note, though singletons (participants who are the sole members of their family cluster) did not contribute to estimation of the random effects variances themselves, they did contribute to the estimation of the total variance, which allows the model to leverage the full ABCD Study^®^ sample.

When including the full ABCD Study^®^ sample, the total variance in the cognitive phenotypes was larger than the total variance in the twin sub-sample. In the full sample, *C* explained a greater proportion of the total variance than in the twin sub-sample where *C* was small for most phenotypes. This result would indicate that the twin sub-sample may be more homogenous compared to the full study sample with respect to common environment. The twin sub-sample consists of twins recruited via birth registries at the four “Twin Hub” sites (Iacono et al. 2018), whereas the full ABCD Study^®^ sample was recruited across twenty one sites, primarily through demographically informed school selection (for details, see Garavan et al. 2018). A study of over 18 million births spanning 72 countries found twin birth to be associated with maternal health conditions, health behaviors, and socioeconomic characteristics (Bhalotra and Clarke 2016). It is possible that these differences result in the observed differences in total variance and variance components. The underlying causes of this discrepancy, though beyond the scope of this paper, are a worthwhile topic for future research.

After expanding the model to include the full sample, we tested the effect of an added random effect of twin status (i.e., “pregnancy ID”). This *T* effect could include any components of the environment that are shared between twins but not among siblings. Examples could include shared uterine environment and prenatal factors, such as gestational age; or the fact that twins experience the same environmental events at exactly the same time. To illustrate this point, a pair of twins might experience a global pandemic at exactly the same age, causing them to experience any effects of the event in similar ways. In contrast, if two siblings are different ages at the time of the event, it might have a different age-dependent effect on each of them (despite the fact that it is occurring as part of their “common environment”). Interestingly, although we found evidence for a *T* effect in matrix reasoning, with a compensatory decrease in the heritability estimate when *T* was included in the model, the estimated *T* effect for the other phenotypes of interest was small to none, as shown in Supplementary Figure 1. One factor that may influence this result is potential overlap between *C* and *T*; in other words, the majority of participants with shared environment are twin pairs, rendering *C* and *T* cross-estimable and hard to fit. Future studies, including those without access to a large twin sample, may still benefit from modeling the *T* effect when there are more similar amounts of twin pairs versus sibling pairs.

We next used the complete sample across multiple timepoints, for a total of over 13,000 observations (Figure 4). Adding the second timepoint led to a substantial decrease in the heritability estimate for height, with a similar increase in the *E* component for height. This may be due to several factors, including possible nonadditive genetic effects (e.g., Silventoinen et al. 2008). Conversely, many of the cognitive phenotypes (with the exception of the Little Man Task and the RAVLT) saw an increase in heritability estimates when modeled across multiple timepoints (Figure 4D). It is possible that this phenomenon is related to the well documented increase in apparent heritability of cognitive traits with age (Davis et al. 2009; Haworth et al. 2010), which may be due in part to the gene × environment correlation (Loughnan et al. 2019). The *S* variance component, representing variance that is unexplained by the other random effects, but which remains stable for a given participant, varied by phenotype. Height had a negligible *S* component, which may be due to the large amount of variance that was already explained by genetic and environmental effects. On the other hand, the NIH Toolbox tasks each had a variance component explained by subject-level variance, indicating that variance in these phenotypes may be relatively more stable for a given participant over time. For these tasks, including *S* in our model allows for better explanation of variance that would otherwise be unexplained in a cross-sectional study. The Little Man Task and the RAVLT did not exhibit subject-specific variance, which may be related to higher noise in these measures as evidenced by the large *E* components for both tasks (.79 and .97 in Model 6, respectively).

Of all the models described in this paper, the model including A, C, S, and E, fit across the full ABCD Study^®^ sample and using all timepoints (Model 6), represented the “most complete” model. However, we employed a series of model comparisons to assess the effect of various study design considerations on the random effects variances. First, we examined the change in our model results when using only kinship-derived genetic relatedness, to approximate a study design in which genetic data are not readily available. We found that, as expected, the model fit was worse in this model, but the parameter estimates were generally similar. Of note, we deliberately used the twin sub-sample data to “assign” relatedness values, meaning that for these analyses the twins recruited through the general population were assumed to have a relatedness value of 0.5 regardless of genetic zygosity. Despite this deliberate attempt to increase the error in our model, estimates remained relatively similar, with inflated estimates for the *T* variance component that seemed to compensate for the induced error in relatedness values (Supplementary Figure 2). These results indicate that when using kinship-derived relatedness, variance that would have been attributed to increased genetic relatedness is “shifted” into the *T* component.

We next tested whether including additional covariates in our pre-residualization step would lead to a change in random effects estimates. In general, residualizing for sociodemographic and genetic ancestry covariates led to a decrease in the total residual variance as well as the common environment (*C*) parameter estimate. This was expected, as adjusting for additional covariates led to a better model fit; the improved model fit was accompanied by a smaller amount of residual variance that was not accounted for by the fixed effects, and any variance that would have been partitioned into *C* was already attributed to the covariates such as household income or parental education. Notably, adjusting for genetic principal components is an attempt to include potential influences of population stratification in the model, and does not influence the estimation of genetic relatedness. Due to the nesting structure of the random effects, the FEMA package only uses the pairwise genetic similarity between individuals within the same family. This contrasts with the genetic principal components, which are used to estimate a fixed effect across the whole sample that may represent population stratification and other effects of genetic ancestry.

Finally, we tested whether omitting the twin sub-sample led to a difference in model results. This analysis aimed to shed light on whether large cohort studies that do not contain a twin-enriched sample can still provide estimates of heritability and other variance components. Overall, the model estimates only slightly changed for most phenotypes, with the exception of the picture sequence memory task which saw a decrease of 0.19 in its heritability estimate. The confidence intervals generated by the two models were similar, suggesting that a large study sample with many siblings is capable of generating model estimates that are similar to those in a twin-enriched sample. Specifically, there were over a thousand families containing more than one participant, leading to several hundred families remaining after the removal of the twin sub-sample. The large number of siblings, perhaps in combination with the use of SNP-derived relatedness covering a range of values, as well as the existence of a smaller number of twins in the sample, were likely related to the similarity between these results and those obtained from the twin sub-sample. Conversely, although large cohort studies such as the ABCD Study^®^ are becoming increasingly common, these results also imply that a smaller twin study may be able to achieve similar heritability estimates to those generated through larger cohort studies.

The results from this study should be considered in light of certain limitations. Generally, LMEs are used to partition the variance in a phenotype of interest into components modeled by random effects; however, models are often built with the assumption that the random effects are mutually independent and follow the normal distributions with mean 0 (Neale and Maes 2004; Wang et al. 2011). Additionally, LMEs represent a “top-down” heritability estimation method that can be biased by several factors including gene–environment correlations, selection, non-random mating, and inbreeding (Zaitlen and Kraft 2012; Zhang and Sun 2022). Furthermore, we did not explore non-additive genetic effects, which can attenuate bias of heritability estimates (Wang et al. 2011); nor did we model any gene × environment interactions, which are likely to exist for some of the phenotypes of interest (Loughnan et al. 2019). Finally, we did not consider sibling interaction in our models, which may interfere with model results due to the potential for different variances in MZ twins compared to DZ twins and full siblings (Eaves 1976).

This work describes many of the modeling techniques available for researchers interested in applying the ACE model and its extensions to a large sample with high relatedness such as the ABCD Study^®^ sample. Notably, the FEMA package provides a tool for mass univariate estimation of LMEs, and its current implementation does not allow for bivariate mixed models. The ability to incorporate multivariate genetic analysis is a major strength of SEM including *OpenMx* (Neale and Maes 2004), and often these techniques are used for more complex models such as factor analysis or interactions among siblings (see Eaves 1976 for an in-depth discussion of sibling interaction). SEM and other implementations of bivariate linear mixed models may therefore provide an avenue to address questions involving genetic and environmental correlations between variables, as well as changes in variance component estimates over time. Bivariate models may also provide some insight into questions of innovation, i.e., whether the set of genes that influence a given phenotype changes over time. In contrast, the mixed effects model used by FEMA leverages the mass univariate approach in a way that allows models to scale to very large datasets spanning multiple timepoints (Fan et al. 2021).

The last several years have seen the development of several new techniques that can be used to model additional relationships, such as random effect × time interaction (He et al. 2016), random effect × covariate interaction (Arbet et al. 2020), covariance among random effects (Zhou et al. 2020; Dolan et al. 2021), and allowing random effects estimates to vary as a function of the phenotype (Azzolini et al. 2022). The sparse clustering design employed in the FEMA package leads to improved computational efficiency compared to other LME implementation software (Fan et al. 2021); future work will investigate the use of FEMA to estimate random effects estimates in more high-dimensional datasets, such as the brain imaging data present in the ABCD Study^®^, and compare with other computationally efficient implementations of the ACE Model such as Accelerated Permutation Inference for the ACE Model (APACE; Chen et al. 2019) and positive semidefinite ACE (PSD-ACE; Risk and Zhu 2021). More broadly, as stated by Zyphur and colleagues (2013), “top down” heritability estimates should serve as just one piece of the puzzle connecting genes and the environment, where current techniques at the molecular and single-gene level may be useful in filling in the gaps from the bottom up.

## Acknowledgements and Conflicts of Interest

The authors wish to thank the youth and families participating in the ABCD Study^®^ and all staff involved in data collection and curation. Dr. Dale reports that he was a Founder of and holds equity in CorTechs Labs, Inc., and serves on its Scientific Advisory Board. He is a member of the Scientific Advisory Board of Human Longevity, Inc. He receives funding through research grants from GE Healthcare to UCSD. The terms of these arrangements have been reviewed by and approved by UCSD in accordance with its conflict of interest policies. Dr. Andreassen has received speakeŕs honorarium from Sunovion, Lundbeck and Janssen, and is a consultant for CorTechs.ai. The remaining authors have no conflicts of interest.

## Ethics approval

The ABCD Study^®^ protocols were approved by the University of California, San Diego Institutional Review Board.

## Consent to Participate

Parent/caregiver permission and child assent were obtained from each participant.

## Consent for Publication

Not applicable.

## Availability of Data and Material

ABCD Study^®^ data release 4.0 is available for approved researchers in NIMH Data Archive (NDA; http://dx.doi.org/10.15154/1523041).

## Code Availability

Code for this project is available at https://github.com/dmysmith/behav-genet-2023. FEMA is available at https://github.com/cmig-research-group/cmig_tools.

## Authors’ contributions

**Conceptualization:** Diana M. Smith, Anders M. Dale; **Methodology:** Diana M. Smith, Robert Loughnan, Naomi P. Friedman, Oleksander Frei, Wesley K. Thompson, Michael Neale, Anders M. Dale; **Formal analysis and investigation:** Diana M. Smith; **Writing - original draft preparation:** Diana M. Smith; **Writing - review and editing:** Robert Loughnan, Naomi P. Friedman, Pravesh Parekh, Wesley K. Thompson, Ole A. Andreassen, Michael Neale, Terry L. Jernigan, Anders M. Dale; **Funding acquisition:** Diana M. Smith, Terry L. Jernigan; **Resources:** Oleksander Frei, Michael Neale, Anders M. Dale; **Supervision:** Wesley K. Thompson, Ole A. Andreassen, Michael Neale, Terry L. Jernigan, Anders M. Dale.

