## Supplementary Information for "Heritability estimation of cognitive phenotypes in the ABCD Study^®^ using mixed models"

**Author Affiliations:**
[1]Neurosciences Graduate Program, University of California San Diego, La Jolla, CA.
[2]Center for Human Development, University of California, San Diego, La Jolla, CA.
[3]Center for Multimodal Imaging and Genetics, University of California, San Diego School of Medicine, La Jolla, CA.
[4]Population Neuroscience and Genetics Lab, University of California, San Diego, La Jolla, CA.
[5]Institute for Behavioral Genetics and Department of Psychology and Neuroscience, University of Colorado Boulder, Boulder, CO.
[6]NORMENT, Division of Mental Health and Addiction, Oslo University Hospital & Institute of Clinical Medicine, University of Oslo, Oslo, Norway
[7]Centre for Bioinformatics, Department of Informatics, University of Oslo, Oslo, Norway
[8]Center for Population Neuroscience and Genetics, Laureate Institute for Brain Research, Tulsa, OK.
[9]Virginia Institute for Psychiatric and Behavioral Genetics, Virginia Commonwealth University, Richmond, VA.
[10]Department of Cognitive Science, University of California, San Diego, La Jolla, CA.
[11]Department of Radiology, University of California, San Diego School of Medicine, La Jolla, CA.
[12]Department of Psychiatry, University of California, San Diego School of Medicine, La Jolla, CA.
[13]Department of Neuroscience, University of California, San Diego School of Medicine, La Jolla, CA.

**Corresponding Author:**
Diana M. Smith
9452 S Medical Ctr Dr
4th Floor, #4W217 C12
La Jolla, CA 92037


### Extended Methods

Below are details of each of the phenotypes analyzed in this study, adapted from Palmer and colleagues (2021b).

***Height.*** We used height data from the baseline visit (mean age = 9.91 years, standard deviation [SD] = 0.63 years) and the year 2 visit (mean age = 11.94 years, SD = 0.65 years). Height was measured as the average of up to 3 separate measures using professional grade equipment (e.g., physician weight beam scale with height rod; Palmer et al. 2021a). After inspection of height data, participants with extreme values (less than 20 inches or greater than 80 inches) were excluded from the sample due to the possibility of errors in data collection and reporting.

***Toolbox Oral Reading Recognition Task® (TORRT):*** measured language decoding and reading. Children were asked to read aloud single letters or words presented in the center of an iPad screen. The research assistant marked pronunciations as correct or incorrect. Extensive training was given prior to administering the test battery. Item difficulty was modulated using computerized adaptive testing (CAT).

***Toolbox Picture Vocabulary Task® (TPVT)***: a variant of the Peabody Picture Vocabulary Test (PPTV), measured language and vocabulary comprehension. Four pictures were presented on an iPad screen as a word was played through the iPad speaker. The child was instructed to point to the picture, which represented the concept, idea or object name heard. CAT was implemented to control for item difficulty and avoid floor or ceiling effects.

***Toolbox Pattern Comparison Processing Speed Test® (TPCPST)***: measured processing speed. Children were shown two images and asked to determine if they were identical or different by touching the appropriate response button on the screen. This test score is the sum of the number of items completed correctly in the time given.

***Toolbox List Sorting Working Memory Test® (TLSWMT)***: measured working memory. Children heard a list of words alongside pictures of each word and were instructed to repeat the list back in order of their actual size from smallest to largest. The list started with only 2 items and a single category (e.g., animals). The number of items increased with each correct answer to a maximum of seven. The child then progressed to the next stage in which two different categories were interleaved. At this stage children were required to report the items back in size order from the first category followed by the second category. Children were always given two opportunities to repeat the list correctly before the experimenter scored the trial as incorrect.

***Toolbox Picture Sequence Memory Test® (TPSMT)***: measured episodic memory. Children were presented with a sequence of pictured objects and activities in a fixed order on a computer screen and simultaneously verbally described, that the participant was asked to remember and then reproduce over three learning trials.

***Toolbox Flanker Task® (TFT):*** measured executive function, attentional and inhibitory control. This adaptation of the Eriksen Flanker task (Eriksen and Eriksen 1974) captures how readily a participant is influenced by the congruency of stimuli surrounding a target. On each trial a target arrow was presented in the center of the iPad screen facing to the left or right and was flanked by two additional arrows on both sides. The surrounding arrows were either facing in the same (congruent) or different (incongruent) direction to the central target arrow. The participant was instructed to push a response button to indicate the direction of the central target arrow. Accuracy and reaction time scores were combined to produce a total score of executive attention, such that higher scores indicate a greater ability to attend to relevant information and inhibit incorrect responses.

***Toolbox Dimensional Change Card Sort Task® (TDCCS)***: measured executive function and cognitive flexibility. On each trial, the participant was presented with two objects at the bottom of the iPad screen and a third object in the middle. The participant was asked to sort the third object by matching it to one of the bottom two objects based on either color or shape. In the first block participants matched based on one dimension and in the second block they switched to the other dimension. In the final block, the sorting dimension alternated between trials pseudorandomly. The total score was calculated based on speed and accuracy.

***Rey-Auditory Verbal Learning Task (RAVLT)***: measures auditory learning, recall and recognition. Participants listened to a list of 15 unrelated words and were asked to immediately recall these after each of five learning trials. A second unrelated list was then presented, and participants were asked to recall as many words as possible from the second list and then recall words again from the initial list. Following a delay of 30 minutes (during which other non-verbal tasks from the cognitive battery are administered), longer-term retention was measured using recall and recognition. This task was administered via an iPad using the Q-interactive platform of Pearson assessments (Daniel et al. 2014). In the current study, the total number of items correctly recalled across the five learning trials was summed to produce a measure of auditory verbal learning.

***Little Man Task (LMT):*** measures visuospatial processing involving mental rotation with varying degrees of difficulty (Acker 1982). A rudimentary male figure holding a briefcase in one hand was presented on an iPad screen. The figure could appear in one of four positions: right side up vs upside down and either facing the participant or with his back to the participant. The briefcase could be in either hand. Participants indicated which hand the briefcase was in using one of two buttons. Performance across the 32 trials was measured by the percentage of trials in which the child responded correctly. This was divided by the average reaction time to complete the task (in seconds) to produce a measure of efficiency of visuospatial processing. This was the dependent variable analyzed in this study.

***WISC-V Matrix reasoning***. Nonverbal reasoning was measured using an automated version of the Matrix Reasoning subtest from the Weschler Intelligence Test for Children-V (WISC-V; Wechsler 2014). On each trial the participant was presented with a series of visuospatial stimuli, which was incomplete. The participant was instructed to select the next stimulus in the sequence from four alternatives. There were 32 possible trials and testing ended when the participant failed three consecutive trials. The total raw score, used in the current study, was the total number of trials completed correctly.

### Supplementary Table 1

| Phenotype | Twin Heritability | Full sample, Baseline | Full Sample, Longitudinal[a] |
|---|---|---|---|
| WISC-V Matrix Reasoning | 0.4 - 0.49 | 0.27 - 0.51 | NA |
| Little Man Task | 0.09 - 0.15 | 0 - 0.09 | 0 - 0 |
| Rey Auditory Verbal Learning Task | 0 - 0 | 0 - 0 | 0 - 0 |
| NIH Toolbox Pattern Comparison | 0.09 - 0.26 | 0 - 0.11 | 0.19 - 0.44 |
| NIH Toolbox Flanker | 0.11 - 0.23 | 0.12 - 0.2 | 0.14 - 0.43 |
| NIH Toolbox Card Sort | 0.32 - 0.39 | 0.24 - 0.33 | NA |
| NIH Toolbox List | 0.25 - 0.34 | 0.17 - 0.27 | NA |
| NIH Toolbox Picture | 0.35 - 0.42 | 0.36 - 0.43 | 0.21 - 0.42 |
| NIH Toolbox Picture Vocabulary | 0.24 - 0.34 | 0.1 - 0.33 | 0.2 - 0.32 |
| NIH Toolbox Reading | 0.56 - 0.58 | 0.41 - 0.57 | 0.5 - 0.63 |
| NIH Toolbox Crystallized Cognition | 0.47 - 0.59 | 0.3 - 0.59 | 0.39 - 0.58 |
| NIH Toolbox Fluid Cognition | 0.48 - 0.55 | 0.24 - 0.49 | NA |
| NIH Toolbox Total Composite Cognition | 0.6 - 0.61 | 0.37 - 0.61 | NA |
| Height | 0.86 - 0.9 | 0.84 - 0.9 | 0.67 - 0.72 |

**Supplementary Table 1.** Summary of heritability estimates. [a]Not all phenotypes were collected at Year 2; only those with available data are shown.

### Supplementary Figure 1

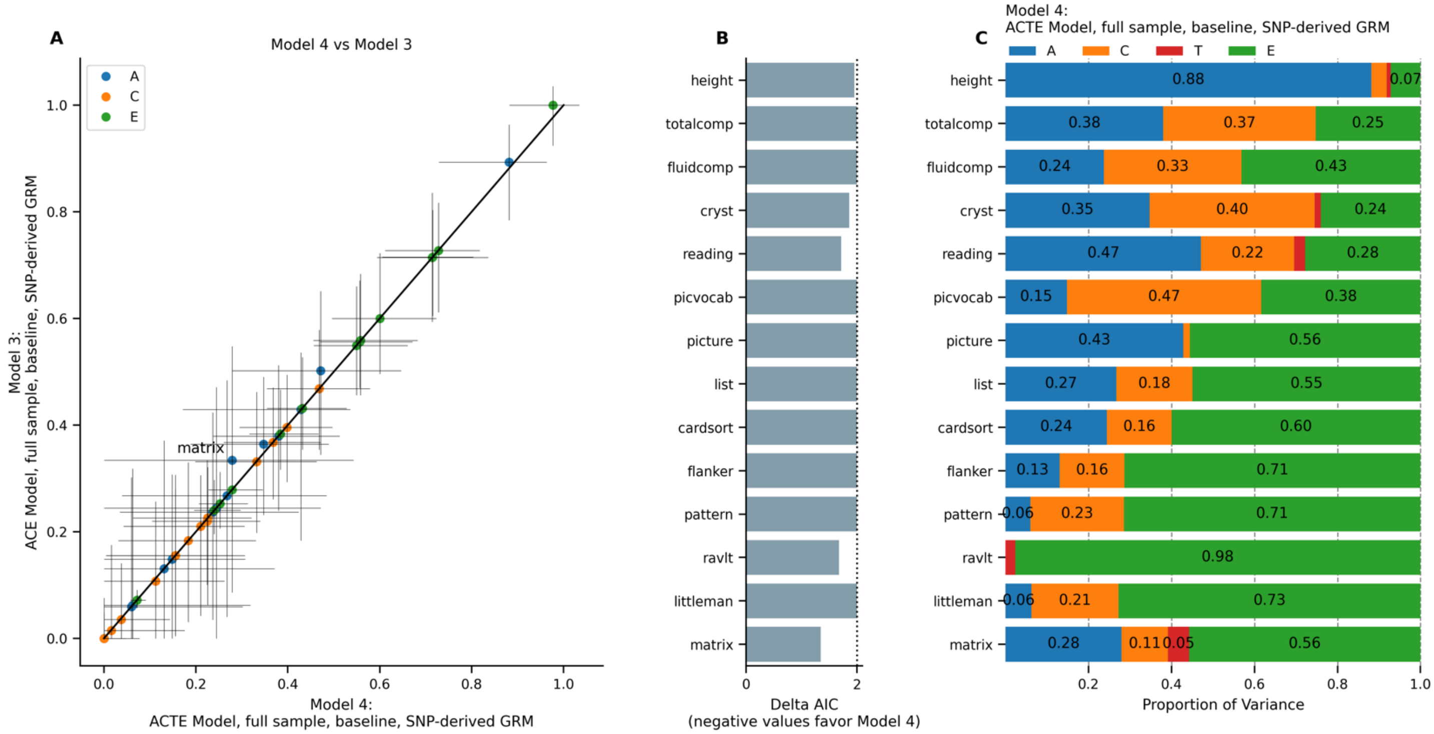

**Supplementary Figure 1.** ACE Model (Model 3) versus ACTE model (Model 4) using full sample at baseline. A) Comparison of model estimates. Horizontal error bars represent confidence interval calculated in Model 4; vertical error bars represent confidence intervals calculated in Model 3. B) Difference in Akaike Information Criterion in Model 4 versus in Model 3. C) Random effects estimates from Model 4.

### Supplementary Figure 2

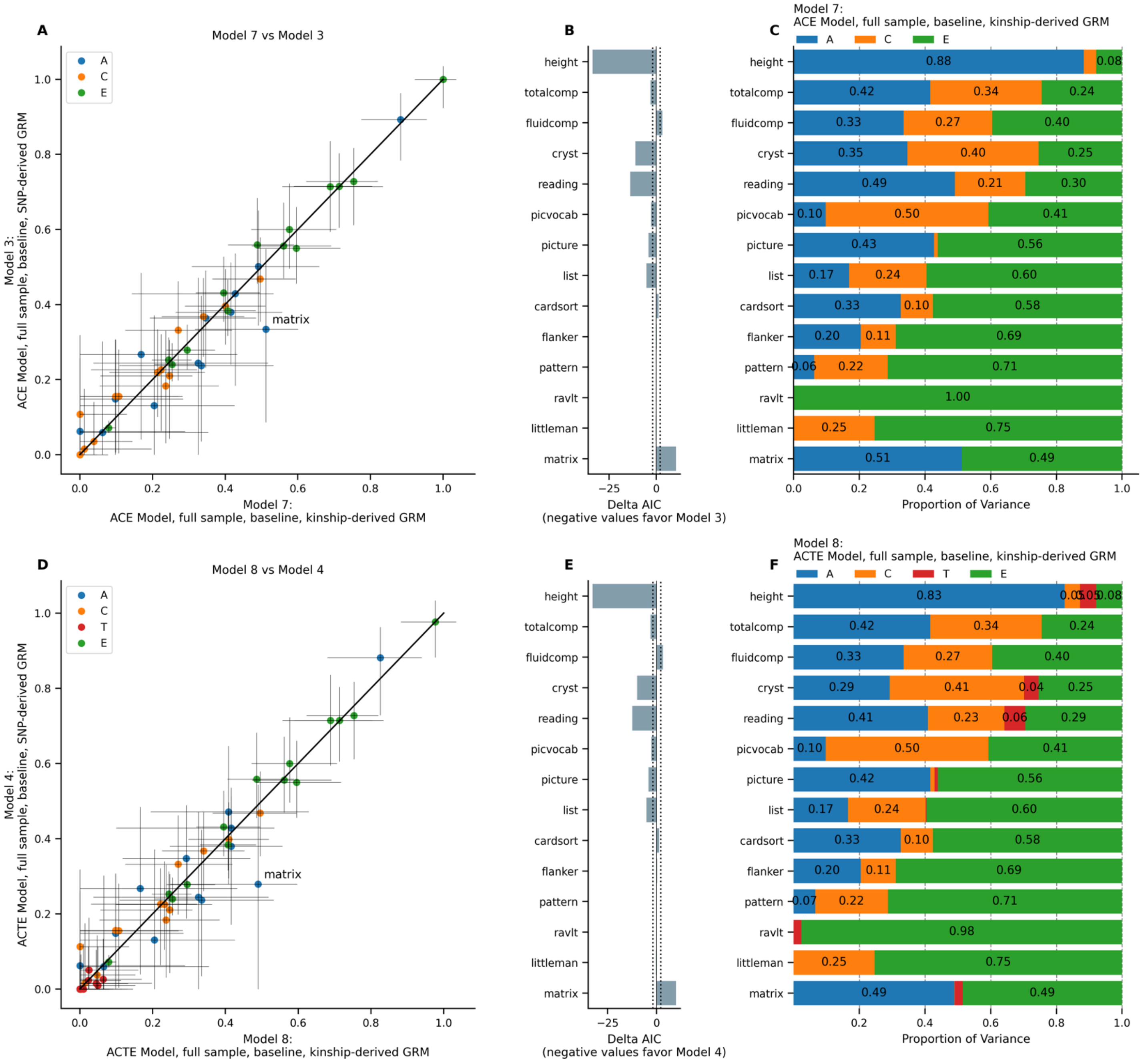

**Supplementary Figure 2.** Model comparison using kinship-derived versus SNP-derived genetic relatedness. A) Comparison of estimates from Model 7 versus Model 3. Horizontal error bars represent confidence interval calculated in Model 7; vertical error bars represent confidence intervals calculated in Model 3. B) Difference in Akaike Information Criterion in Model 7 versus in Model 3. C) Random effects estimates from Model 7. D) Comparison of estimates from Model 8 versus Model 4. Horizontal error bars represent confidence intervals calculated in Model 8; vertical error bars represent confidence intervals calculated in Model 4. E) Difference in Akaike Information Criterion in Model 8 versus in Model 4. F) Random effects estimates from Model 8.

### Supplementary Figure 3

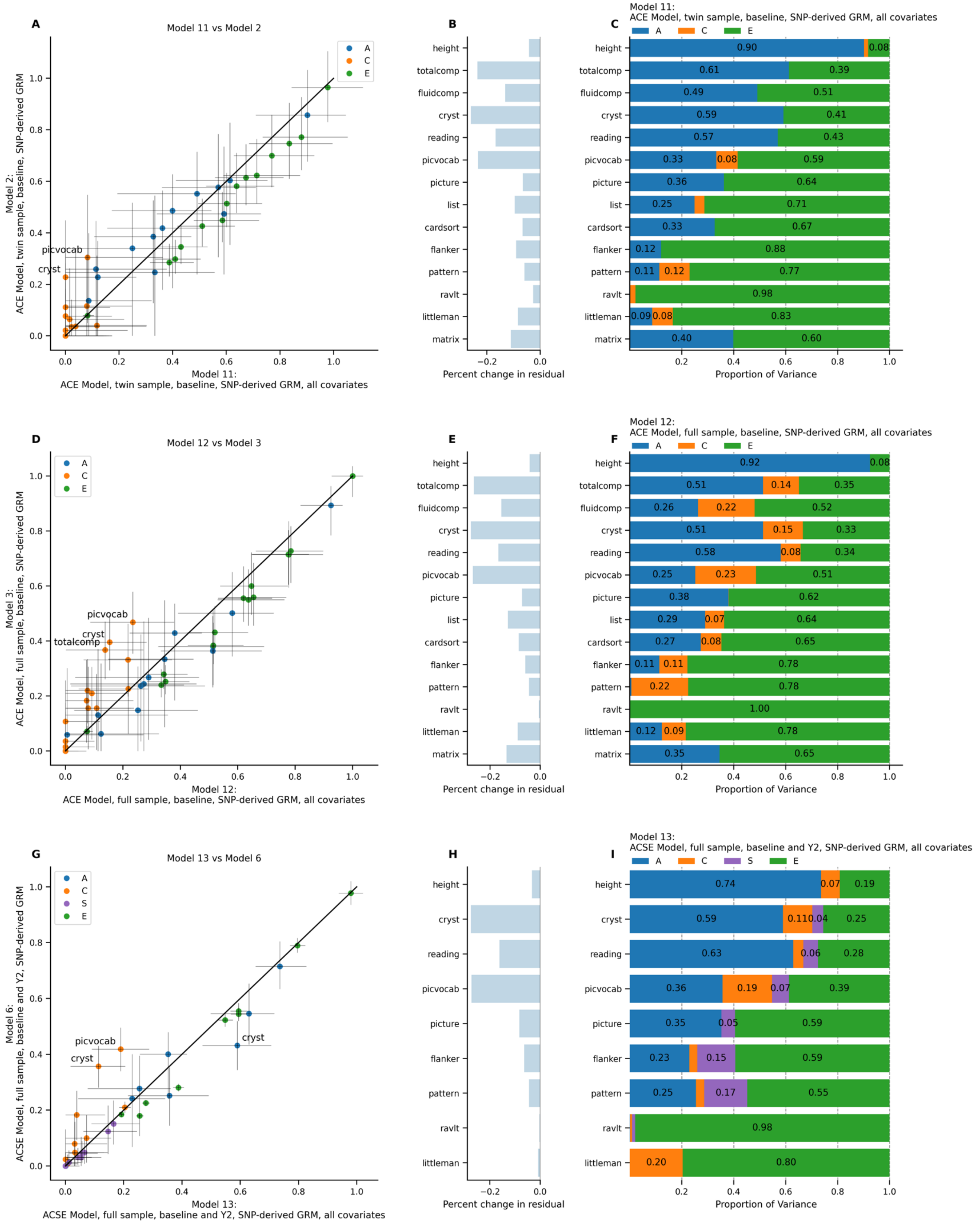

**Supplementary Figure 3.** Model comparison after residualizing for all covariates versus age and sex only. A) Comparison of estimates from Model 11 versus Model 2. Horizontal error bars represent confidence interval calculated in Model 11; vertical error bars represent confidence intervals calculated in Model 2. B) Difference in total residual variance in Model 11 versus in Model 2. C) Random effects estimates from Model 11. D) Comparison of estimates from Model 12 versus Model 3. Horizontal error bars represent confidence intervals calculated in Model 12; vertical error bars represent confidence intervals calculated in Model 3. E) Difference in total residual variance in Model 12 versus in Model 3. F) Random effects estimates from Model 12. G) Comparison of estimates from Model 13 versus Model 6. Horizontal error bars represent confidence intervals calculated in Model 13; vertical error bars represent confidence intervals calculated in Model 6. H) Difference in total residual variance in Model 13 versus in Model 6. I) Random effects estimates from Model 13.
